# HIV-1 gp120 impairs spatial memory through CREB

**DOI:** 10.1101/2020.10.20.347146

**Authors:** Jenny Shrestha, Maryline Santerre, Charles NS Allen, Sterling P Arjona, Ruma Mukerjee, Jin Park, Asen Bagashev, Ying Wang, Marcus Kaul, Jeannie Chin, Bassel E Sawaya

**Author notes:** **Corresponding Author:** Bassel E Sawaya, PhD, MS(IME), MBA, Fels Institute for Cancer Research, Lewis Katz School of Medicine, Temple University. 3307 North Broad Street; Philadelphia, Pennsylvania 19140. JS-Bristol Myers Squibb, 556 Morris Ave (Building S7), Summit, NJ 07901 AB-Department of Cancer Pathobiology, The Children’s Hospital of Philadelphia, 4300 Colket Translational Research Bldg; 3501 Civic Center Blvd, Philadelphia, PA 19104.

## Abstract

HIV-associated neurocognitive disorders (HAND) remains an unsolved problem in the clinical management of HIV-1 carriers, because existing anti-retroviral therapy while suppressing viral replication, do not prevent neurocognitive impairment (e.g. spatial memory loss). HIV-1 gp120 protein has been proposed to contribute to HAND because it is shed by infected cells and the use of antibodies revealed its presence in cerebrospinal fluid (CSF) even in the combinatory antiretroviral therapy (cART) era. The cyclic AMP response element-binding protein (CREB) has long been known to be a star player in memory. CREB exerts its effect partially through regulating the genes for peroxisome proliferator-activated receptor gamma coactivator (PGC)-1α and brain-derived neurotrophic factor (BDNF). CREB, PGC-1α, and BDNF levels are low in the brains of patients with neurodegenerative diseases and a dearth of either protein is associated with cognitive decline. We have obtained data showing that gp120 contributes to neurodegeneration by altering CREB phosphorylation on serine residue 133 thus disrupting mitochondrial movement and synaptic plasticity leading to spatial memory loss. Inhibition of CREB function was also associated with a decrease of ATP levels and lower mitochondrial DNA copy numbers. Our data was validated *in vitro* (primary mouse neurons and neuronal cell line, SH-SY5Y) and *in vivo* (gp120-tg mice and mice injected with gp120). The negative effect of gp120 was alleviated in cells and animals in the presence of Rolipram. Hence, we conclude that HIV-1 gp120 protein contributes to spatial memory impairment via inhibition of CREB protein activity.

## INTRODUCTION

Patients infected with HIV-1, including those using the effective but not curative combinatory antiretroviral therapy (cART) suffer from deregulation and impairment of organs such as the heart, kidney, and brain (1). Studies involving 37,000 HIV-infected patients using cART (from 2003-2013) showed that comorbidity increased with age in those patients compared to uninfected age-matched patients. Similarly, a longitudinal study involving over 5,000 patients confirmed this observation and found an unequivocal link between HIV-1 infection and premature brain aging (2).

The persistence of neurocognitive disorders could be due to (i) the limitation and toxicity of cART, (ii) the reactivation of latent viruses, (iii) inflammation caused by secreted toxins (e.g. TNF-α), (iv) substance of abuse (drugs, alcohol, tobacco), and/or (v) the presence of “defective proviruses” that are capable of transcribing novel translationally competent unspliced HIV-RNA species. The latter explains the detection of viral proteins such as Tat and gp120 in cerebrospinal fluid (CSF) (3). These defective proviruses are not “silent”, as they contribute to HIV-1 pathogenesis independently of cART (4, 5) and control viral latency. As a result of this persistence, a significant number of HIV-1 patients suffer from cognitive impairments such as spatial memory loss.

HIV-1 gp120 is an envelope protein that allows the interaction of the virus with the host receptors (CD4^+^) and co-receptors (CCR5 and CXCR4). HIV-1 gp120 is released from HIV-infected cells and can cause neuronal dysfunction (6). gp120 was reported to be toxic to neurons through activation of the NMDA receptor, causing an increase in Ca^2+^ influx, activating the oxidative stress (OS) pathway, and allowing the release of toxic lipids from membranes (7). gp120 protein was also shown to alter mitochondrial functions (8) and axonal transport (9) causing neuronal deregulation. Studies showed that knockout and/or knockdown of CCR5 protects against gp120 V3 peptide-induced memory deficits (10).

*Per* the literature, studies unequivocally demonstrated the cAMP response element-binding protein (CREB) is the key player in functional memory and that its dephosphorylation on Serine 133 leads to hippocampal episodic memory impairments (11,12). In healthy brains, synaptic activity induces or represses specific genes, which in turn leads to the long-lasting synaptic changes that underlie learning and memory. CREB protein plays a crucial role, in part, by regulating the gene for brain-derived neurotrophic factor (BDNF), a powerful neuronal growth, and neuroprotective protein with numerous functions in the brain. CREB and BDNF levels are low in the brains of people with Alzheimer’s disease (AD) and other neurodegenerative diseases and a dearth of CREB and/or BDNF is associated with cognitive decline (13).

CREB protein was first described as a cAMP-responsive transcription factor regulating the somatostatin gene (14). It binds to DNA sequences called *cAMP response elements* (CRE), thereby increasing the transcription of the downstream genes such as BDNF, somatostatin, Bcl-2, c-fos, proliferator-activated receptor gamma coactivator (PGC)-1α, and interleukin (IL)-10.

CREB can be phosphorylated on various residues, however, its phosphorylation on serine residue 133 by protein kinase A (PKA) and other kinases is critical for binding to CRE motifs (15). CREB plays a role in the regulation of short- and long-term memory formation (16). It is essential to the formation of spatial memory (17) and plays a positive role in axonal transport and synaptic plasticity (18,19). CREB protects the neurons through the regulation of the PGC-1α, Bcl-2, and BDNF promoters. In the adult brain, CREB is involved in learning, memory, and neuronal plasticity. BDNF is the main gene important for synaptic plasticity, memory consolidation, and long-term potentiation and is targeted by CREB (20). Therefore, it is expected that inhibition of CREB protein will have multiple damaging outcomes contributing to spatial memory impairment observed in patients infected with HIV-1 and suffering from HAND (21).

The relationship between CREB and spatial memory is well established. For example, decreased expression of pCREB^S133^ in the hippocampus and amygdala of aged rats was described to contribute to the rapid loss of spatial memory as tested by inhibitory avoidance training (22).

Decreased pCREB^S133^ expression was detected in the cerebral cortex and hippocampus of aged rats suffering from memory loss (23). Low expression of pCREB^S133^ was shown to be associated with spatial memory impairment in the hippocampus of old rats (120 weeks) compared to young rats (15 weeks) (24). Similarly, aged mice subjected to contextual fear assay expressed less pCREB^S133^ and increased spatial memory impairment. Another study showed somatic gene transfer of CREB protein attenuates spatial memory impairment in 15 months old rats (25).

Finally, an increase of pCREB^S133^ expression was shown in old mice that received the blood of young mice, and significant progress in their spatial memory was also observed (26). The same study also showed that pCREB^S133^ is one of the main contributors to restoring the memory in transgenic mice modeling AD, when receiving young blood plasma.

Therefore, based on memory impairment observed in HIV-1 infected patients in the cART era, the presence of gp120 protein in CSF, and the key role played by phosphorylated CREB protein in spatial memory, all provided the rationale to determine whether gp120 impairs spatial memory through CREB?

## RESULTS

### CREB expression is altered with memory impairment in mice and humans

Reduced CREB expression and loss of its phosphorylation has been observed in patients suffering from diseases, including AD, PD, ALS, and HD. To validate this observation in an AD model, 4-month old APP-transgenic (APP-tg) mouse brains and age-matched non-transgenic littermates were processed and assessed for expression levels of total CREB using immunofluorescence (IF). There appears to be a dramatic decrease in CREB expression in the APP-tg mice compared to the control littermates (Fig 1A).

**Figure 1.**
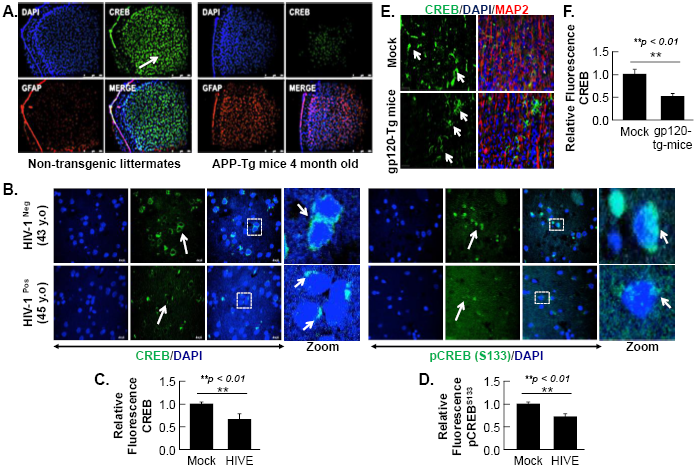
CREB expression level is altered in brain tissues. ***A***. Expression level of CREB protein (green) was observed in the brain tissues of 4-month-old APP-transgenic mice compared to non-transgenic littermates. Immunohistochemistry was performed using antibodies for CREB (green), and glial fibrillary acidic protein (GFAP) to visualize astrocytes (red). Nuclei were stained using DAPI (blue). ***B***. Distribution and expression of total CREB and pCREB^S133^ (green) by immunohistochemistry in the brain tissue (frontal lobe) of a 43-year-old HIV-negative patient and a 45-year-old patient with HIV-1 encephalitis (HIVE). Nuclei were stained using DAPI (blue). ***C., D***. Quantification of the fluorescence intensity of CREB and pCREB^S133^ labeling shown in B. ***E***. The expression of total CREB (green), and MAP2 (red) to visualize neurons, in hippocampal brain slides of 9-month-old HIV-1 gp120 transgenic mice (gp120-Tg) and age matched control mice. Nuclei were stained using DAPI (blue). ***F***. Quantification of the fluorescent intensity of CREB labeling shown in ***E***. Data from panels ***C, D*** and ***F*** represent the mean ± S.D. Results were judged statistically significant if *p*<0.05 by analysis of variance. (\**p*<0.05; \*\**p*<0.01; \*\*\**p*<0.001). Arrows in panels ***A, B*** and ***E*** are pointing at CREB protein.

Next, the expression levels of total CREB and phosphorylated CREB (pCREB^S133^) in the frontal lobe brain tissues of an HIV-infected patient suffering from memory impairment were measured. Total CREB and pCREB^S133^ were shown to be decreased compared to an age-matched uninfected control patient (Fig 1B). Interestingly, CREB expression seems to be perinuclear in both patients, while pCREB^S133^ was partially nuclear in the uninfected mock but not in the HIV-infected patient (panel B, zoom).

Using human brain tissue samples provided by NNTC, we showed that the loss of total and phosphorylated CREB (on serine residue 133) in the brain tissues of HIV-1 infected patients is similar to patients enduring neurodegenerative diseases, and memory impairments (27, 28).

### gp120 alone alters CREB expression in mice in vivo

gp120-tg mice were generated by inserting the portion of the HIV-*env* gene that encodes gp120 into the mouse genome under the control of glial fibrillary acidic protein (GFAP) promoter, resulting in gp120 protein expression specifically in astrocytes (29, 30). We found that the expression level of CREB is altered in the hippocampus of these 9-month old mice (Fig 1E). CREB appeared to be decreased in the hippocampus of these mice compared to the mock suggesting that gp120 alone can result in similar effects to what is seen in HIV-infected patients.

### Recombinant HIV-1 gp120 protein alters CREB expression in primary cell cultures and neuronal cell lines

The neuroblastoma cell line, SH-SY5Y, was used to assess the concentration of gp120 necessary to cause effects on CREB expression and phosphorylation without hindering cell viability severely. SH-SY5Y cells were differentiated with retinoic acid for 72 hours and then treated with various concentrations of recombinant gp120 protein (HIV_IIIB_ gp120, NIH-AIDS Reagents Program) for 24 hours. An MTT assay for cell viability was performed and showed that a 100ng/ml of HIV-1 gp120 protein modestly affects cell viability and enough to dephosphorylate CREB protein. (Fig 2A).

**Figure 2.**
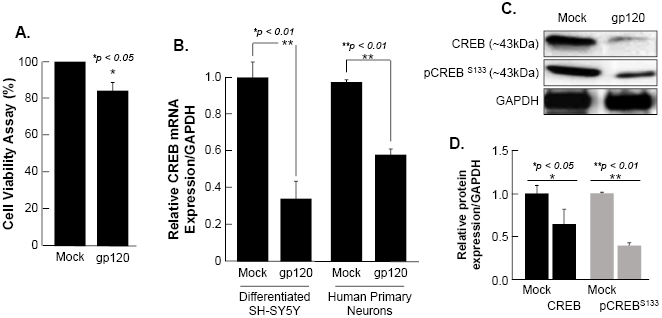
Deregulation of CREB in HIV-1 gp120 treated cells. ***A***. Differentiated SH-SY5Y cells were treated with gp120 protein for 24 hours then subjected to MTT assay to measure cell viability as indicated. The results are statistically significant using Student’s *t*-test. ***B***. Expression of CREB mRNA isolated from differentiated SH-SY5Y and human primary neurons from a 26-week-old fetal brain. These were then treated with 100 ng/ml of HIV-1 gp120 IIIB for 24 hours and CREB mRNA expression was measured by qPCR (repeated 3x) and normalized to GAPDH as an internal control. Data represents the mean ± S.D. Results were judged statistically significant if *p*<0.05 by one-way analysis of variance and ANOVA test. (\**p*<0.05; \*\**p*<0.01; \*\*\**p*<0.001). ***C***. Expression of CREB protein using 50 μg of extracts isolated from untreated or HIV-1 gp120-treated SH-SY5Y as obtained by western blot using anti-CREB, -pCREB^S133^ or -GAPDH (used as protein loading control) antibodies. ***D***. Quantification of the relative protein level was determined from the band intensity using ImageJ software and normalized relative to the GAPDH. Bar graph represent means ± SD of at least two independent experiments.

Next, SH-SY5Y and primary human neurons were treated with 100 ng/ml of recombinant gp120 IIIB protein for 24 hours. mRNA and protein expression levels were then measured by qPCR (Fig 2B). The expression of CREB mRNA is significantly lower in cells treated with HIV-1 gp120 compared to controls. CREB protein (total and phosphorylated) expression levels were also assessed by Western blot and show decreased expression in gp120-treated SH-SY5Y cells (Fig 2C). Changes in CREB expression levels were measured using Image J and presented as histograms (Fig 2D). Next, we sought to determine whether gp120 alters CREB function(s).

### HIV-1 gp120 alters PGC-1α, a protein involved in mitochondrial biogenesis

Among its many functions, CREB protein has been shown to modulate mitochondrial biogenesis, directly and indirectly, through regulating the transcription of several genes such as *pgc-1*α (directly) and *bdnf* (indirectly) (31, 32).

To assess the direct pathway, differentiated SH-SY5Y cells were treated with 100 ng/ml of HIV-1 gp120 for 48 hours. The cells were then divided into two groups where the mRNA was isolated from one group and subjected to qPCR (Fig 3A) while protein extracts were isolated from the second group and processed for Western blot analysis (Fig 3B). As shown, the addition of HIV-1 gp120 protein led to decreased mRNA and protein expression of PGC-1α. Change in PGC-1α protein expression was measured and presented as a histogram in panel B. These data suggest a loss of CREB expression by HIV-1 gp120 leads to loss of PGC-1α expression which could cause downstream deleterious effects on mitochondrial biogenesis.

**Figure 3.**
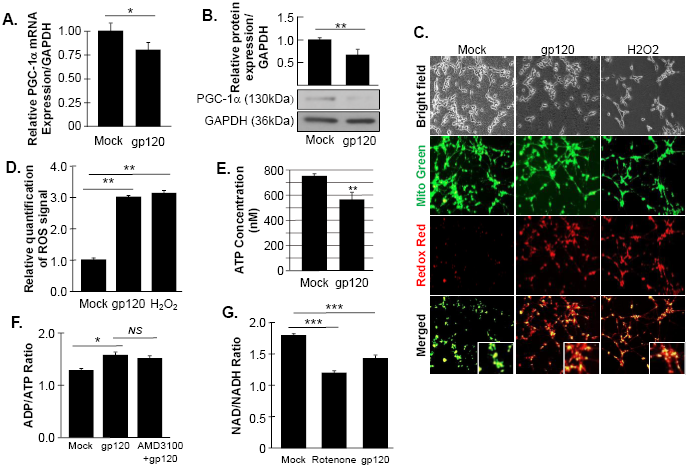
gp120 alters mitochondrial energy. Expression of PGC-1α mRNA and protein isolated from untreated or HIV-1 gp120-treated SH-SY5Y as obtained by qPCR (***A***.) and western blot (***B***.) assays, respectively. Quantitative analysis of the western blot presented along with the bands as a bar graph. ***C***. Differentiated SH-SY5Y cells were treated with 100ng/ml of HIV-1 gp120 for 24 hours or H_2_O_2_ then stained with Redox Sensor Red CC-1 (red) and MitoTracker Green (green). ROS production is seen by co-localization of the two dyes in the mitochondria. Data shown here are from a single that was replicated 3 times and statistically significant. ***D***. Quantification of ROS signal in panel ***C*** using the Student’s *t*-test statistical significance level, *p*<0.05 when compared to the mock control group (one-way ANOVA). ***E***. Concentration of ATP produced by SH-SY5Y cells treated with HIV-1 gp120 compared to mock (untreated) cells. ***F***. Measurement of ADP/ATP ratio in mock, HIV-1 gp120-treated, and AMD3100 + HIV-1 gp120-treated SH-SY5Y cells. AMD3100 (1 μM) (inhibitor of CXCR4) pre-treatment was used as a control to inhibit gp120-CXCR4 association. ***G***. Measurement of NAD/NADH ratio in mock, HIV-1 gp120-treated and rotenone-treated cells. Rotenone is a complex I inhibitor and was used as a positive control. The experiments were done in triplicate and the *p*-values were calculated using Student *t*-test (Mean ± S.D.). (\**p*<0.05; \*\**p*<0.01; \*\*\**p*<0.001).

### HIV-gp120 deregulates mitochondrial energy

The reduction of PGC-1α protein expression sensitizes the cells to oxidative stress, resulting in ROS accumulation and a decrease in ATP generation (33). Therefore, we sought to determine whether the addition of gp120 promotes these events. SH-SY5Y cells were treated with 100 ng/ml of recombinant gp120 or H_2_O_2_ for 24 hours after which the cells were collected and ROS production was measured (Fig 3C). The cells were then stained with Redox Red and Mito Green to visualize oxidation of the cells and the mitochondria, respectively. The Redox Red dye accumulated in the mitochondria of cells treated with HIV-1 gp120 and H_2_O_2_ indicating a high redox potential of the cytosol in these cells (evidenced by the co-localization of the Redox Red and Mito Green dyes). Quantification of ROS is presented in panel D.

Next, we measured the ATP levels in SH-SY5Y cells treated with 100ng/ml of recombinant HIV-1 gp120 protein for 24 hours (Fig 3E). The addition of HIV-1 gp120 decreased the ATP concentration compared to mock cells. To validate these results, the ADP/ATP ratio was also measured (Fig 3F). As expected, the ratio of ADP/ATP increased in HIV-1 gp120-treated cells confirming our previous data (*AMD3100 pre-treatment was used as an inhibitor for CXCR4 to inhibit the effect of gp120*).

To further validate our data, we measured the ratio of NAD^+^/NADH (Fig 3G). Nicotinamide adenine dinucleotide (NAD) is used as a marker for aging and may help to explain the ATP reduction and ROS production seen in these cells. The addition of recombinant HIV-1 gp120 to SH-SY5Y cells lowered the NAD^+^/NADH ratio. Rotenone was used as a positive control because it works by interfering with the electron transport chain in mitochondria. (34) It interferes with NADH during the creation of usable cellular energy (ATP) that can damage DNA and other components of the mitochondria.

### HIV-1 gp120 causes a loss of mitochondrial cristae and alters mitochondrial ultrastructure

The inner membrane of the mitochondria, also known as mitochondrial cristae due to their wavy shape, is the site of electron exchange and ATP production. Loss of the cristae leads to dissociation of ATP synthase dimers and impaired ability of the mitochondria to supply the cell with enough ATP required for cell function (35). SH-SY5Y cells were treated with 100 ng/ml of recombinant HIV-1 gp120 protein for 24 hours and subjected to electron microscopy. The ultrastructure of the mitochondria, including the cristae, was examined and compared to that of control (mock) cells (Fig 4A). the mitochondria shape changed and the cristae were lost in gp120-treated cells. This ultrastructural alteration could explain the decrease in ATP production and increased ROS seen in these cells. No significant changes in the mitochondrial area in (Fig 4B) or perimeter were observed (data not shown).

**Figure 4.**
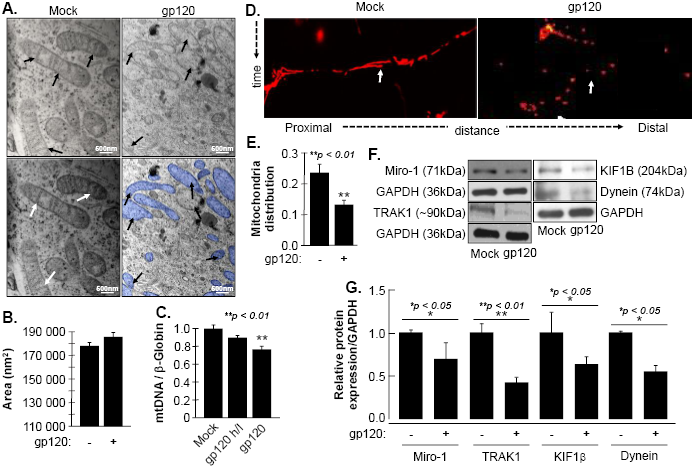
Deregulation of mitochondrial ultrastructure, mtDNA, and distribution in the presence of HIV-1 gp120. ***A***. Differentiated SH-SY5Y cells were treated with either PBS or recombinant HIV-1 gp120 protein at 100ng/ml for 24 hours before being fixed and processed for electron microscopy. Mock cells show mitochondria (highlighted in blue) with intact cristae (arrows) while in HIV-1 gp120-treated cells show fewer and shorter cristae as well as large open spaces (swelling mitochondria) where no visible cristae are observed (black and white arrows). ***B***. Mitochondrial areas were measured (nm^2^) using ImageJ. Any structure that cannot be clearly identified as a mitochondrion (i.e. no evidence of cristae or double membrane) or that has been cut off when acquiring the image was not included. Standard error was used. ***C***. mtDNA copy number was determined by taking the mtDNA/β-Globin ratio. Fold change in mtDNA copy number in HIV-1 gp120-treated SH-SY5Y cells relative to the mock or heat inactivated (h/I) HIV-1 gp120 protein (protein was heated at 90°C for 10 minutes) is shown. Bars represent mean values ± S.E.M. p<0.001. ***D***. A representation of the mitochondria distribution, and density in the neuronal processes of the differentiated SH-SY5Y cells. Images were taken after transfecting the cells with 0.5 μg of Mito dsRED plasmid prior to differentiation. The images were taken post 24 hours of the respective treatment using a confocal microscope. Arrows represent the gaps between the mitochondria. ***E***. Ratio of moving mitochondria was measured and represented as a bar graph using the Student’s *t*-test. Statistical significance level, *p*<0.01. ***F***. Expression of motor and mitochondrial proteins (Miro-1, Trak1, KIF1β, and Dynein) isolated from untreated or HIV-1 gp120-treated SH-SY5Y as obtained by western blot analysis. ***G***. Quantitative analysis of the western blot presented along with the bands as a bar graph using the Student’s *t*-test statistical significance level, *p*<0.05 or 0.01.

Low ATP level is also associated with decreased mitochondrial DNA (mtDNA) copy number. Therefore, the mtDNA copy number was measured in the absence and presence of HIV-1 gp120 protein in human primary cultures of neurons (hippocampal) (Figure 4C). The addition of HIV-1 gp120 for 24 hours led to a significantly reduced mtDNA copy number. Note that a 20% change in the mtDNA copy number is considered significant (36).

Next, we examined mitochondrial distribution and density in SH-SY5Y cells treated with 100 ng/ml of recombinant gp120 protein. Untreated cells were used as control. These cells also expressed a Mito dsRed plasmid to visualize the mitochondria. Mitochondria distribution and density were reduced in HIV-1 gp120-treated cells was observed compared to mock untreated as obtained by confocal microscopy and measured using (Fig 4D and 4E).

Additionally, we tested the expression level of Miro-1 (aka Rho T1), TRAK1 (aka Milton), KIF1β, and Dynein proteins. Miro is a protein that links mitochondria to KIF5B and Dynein motor proteins, allowing the mitochondria to move along the microtubule, while TRAK1 is involved in mitochondrial motility. KIF1β and Dynein proteins are involved in the mitochondria movement. The addition of gp120 resulted in a decrease in Miro-1, TRAK1, KIF1β, and Dynein expression levels (Fig 4F). These results confirm the ability of gp120 to cause mitochondrial movement impairments. Changes in protein expression levels were quantified using Image J and presented as histograms (Fig 4G). These results confirm our hypothesis that gp120 causes mitochondrial damage, and suggest it does this through a pathway that involves regulating CREB expression, subsequent regulation PGC-1α, loss of cristae, and subsequent decrease in ATP production and impaired mitochondrial movement.

### Loss of pCREB and PGC-1α cause synaptic plasticity deregulation

It has been described that the loss of PGC-1α reduces the density of dendritic spines in the hippocampus *in vivo* (37). The loss of PGC-1α can also inhibit BDNF-induced dendritic spine formation. Interestingly, BDNF increases the expression of PGC-1α through the activation of CREB protein (38). These observations suggest that a feedback loop exists between the three proteins and affecting one protein might alter the functions of the two others. To that end, we examined the expression level of BDNF protein in the presence of gp120. SH-SY5Y cells were treated with 100 ng/ml of recombinant gp120 for 24 hours after which the cells were collected and processed for qPCR and Western blot analysis (Figure 5A and B). The addition of gp120 protein resulted in a decrease in BDNF mRNA and protein expression levels compared to the mock untreated. Changes in BDNF protein expression levels were measured using Image J (Fig 5B, histogram). These results corroborate previous data regarding the decrease of BDNF in HIV-human brain tissues (39) and the ability of CREB to regulate the BDNF gene. These results also suggest a feedback loop between CREB-BDNF and PGC-1α.

**Fig 5.**
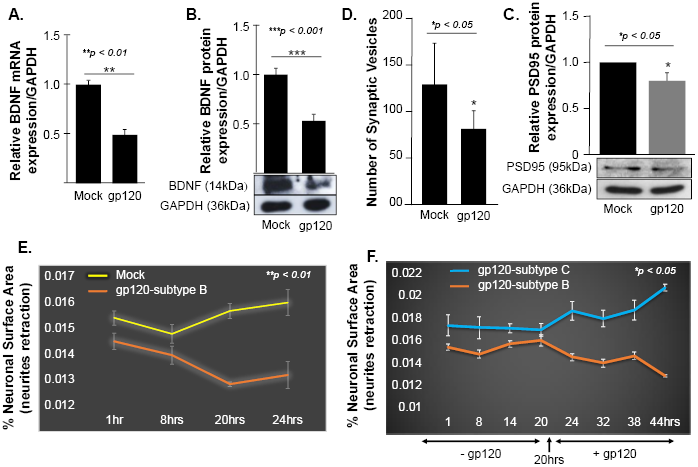
HIV-1 gp120 alters synaptic plasticity. Expression of BDNF mRNA and protein isolated from untreated or HIV-1 gp120-treated SH-SY5Y as obtained by qPCR (***A***.) and Western blot (***B***.), respectively. Quantitative analysis of the western blot presented on the top of the bands as a bar graph. Student’s *t*-test statistical significance level, *p*<0.001. ***C***. *Quantification of synaptophysin-positive synaptic vesicles--* Synaptophysin was expressed using Cell Light Transduction at MOI of 5. HIV-1 gp120 protein or mock treatment was added to primary mouse neurons 24 hours after the transduction. Synaptophysin vesicles were counted using ImageJ for each condition. Images from at least 10 different fields were taken for statistical analysis *p*<0.05. ***D***. Expression of PSD95 protein isolated from untreated or HIV-1 gp120-treated SH-SY5Y as obtained by western blot assays. Quantitative analysis of the western blot presented on the top of the bands as a bar graph. Student’s *t*-test statistical significance level, *p*<0.05. ***E., F***. SH-SY5Y cells were treated with 100ng/ml of HIV-1 gp120 (subtype B) (***E***.). Live cell images and bright field contract images were acquired every 30 minutes from the same field for the indicated times. The images were analyzed with ImageJ software and the surface area covered by the cells was normalized against the total number of the cells in each time point. The experiment was repeated with gp120 (clades B and C) (***F***.) Supplemented medium added at 20 hours. The experiments were repeated 3 times and the images from at least 10 different fields were taken for each set for the statistical significance (*p*<0.05.).

It has been shown that mitochondria, CREB, and BDNF play a significant role in controlling fundamental processes in neuroplasticity (40). Further, reduction or absence of CREB phosphorylation affects synaptic plasticity that is important for memory and has been linked to long-term potentiation and PSD-95 regulation (41). Therefore, we determined the effect of CREB-BDNF and PGC1a loss on two synaptic markers, synaptophysin (a protein involved in synaptic transmission) and PSD-95 (is a scaffold protein present on the dendrites of the postsynaptic neurons and has been used as a synaptic marker, and stabilization of synaptic changes during long term potentiation).

Primary mouse neurons (E18) were isolated from the hippocampal area, cultured and their synaptophysin vesicles were labeled with red fluorescence (Thermo Fischer) for 22 hours before adding 100 ng/ml of recombinant gp120 protein for an additional 24 hours (Fig 5C). Images were acquired on live cells 24 hours post gp120 treatment using the EVOS AMD microscope. We observed a decrease in the number of synaptophysin vesicles in gp120-treated cells compared to the mock untreated cells.

Next, we examined the expression level of PSD-95. Differentiated with 10 μM of retinoic acid, SH-SY5Y cells were treated with 100 ng/ml of recombinant gp120 for 24 hours. The cells were collected, protein extracts were isolated and subjected to Western blot analysis using anti-PSD-95 or -GAPDH antibodies. The expression level of PSD-95 protein decreased in HIV-1 gp120-treated cells compared to the mock untreated cells (Fig 5D).

Low expression of PSD-95 protein points to synapse shortening and loss of synaptic plasticity. We validated this observation by measuring the neuronal surface area in live cells as an evaluation of neurite retraction and thus synapse shortening (Fig 5E). The shortening in neurite length was observed in SH-SY5Y cells treated with HIV-1 gp120 compared to mock cells.

To further validate these results, we repeated this experiment using 100 ng/ml of recombinant gp120 proteins prepared from HIV-1 (clades B or C). We found that the addition of gp120-B but not gp120-C causes shortening of neurites length in SH-SY5Y (Fig 5F). HIV-1 gp120 (subtype C) was shown to be responsible for decreased neurovirulence of clade C. These results suggest that HIV-1 gp120 can cause neurite retraction *in vitro* possibly leading to altered spatial memory *in vivo*.

### HIV-1 gp120-induced mitochondrial dysfunction is CREB-dependent

So far, our data has shown HIV-1 gp120 has the ability to promote dephosphorylation of CREB leading to detrimental downstream effects. Thus, we wanted to see whether we can restore CREB phosphorylation and function (using mtDNA copy number, ATP production, mitochondrial movement, and oxygen consumption rate as readouts) by using a selective PDE4 inhibitor, rolipram.

Rolipram (C_16_H_21_NO_3_) was developed as a potential antidepressant drug in the early 1990s (42, 43). Many functions were associated with Rolipram use in animals including restoring spatial memory through reactivation of CREB. Rolipram has since been discontinued due to its significant side effects.

Ethanol (EtOH), a proven CREB inhibitor and known to cause short memory lapse by dephosphorylating CREB protein and inhibiting its function (44, 45) was used as a negative control. HEK-293 cells, cultured in duplicate, were treated with 200 mM of EtOH for 24 hours or with 30 μM of Rolipram for 1 hour. The cells were collected, the first set was subjected to Western blot analysis using anti-total CREB, -pCREB^S133^, and -GAPDH antibodies, while mitochondrial DNA (mtDNA) copy number was measured in the second set (Fig 6A).

**Fig 6.**
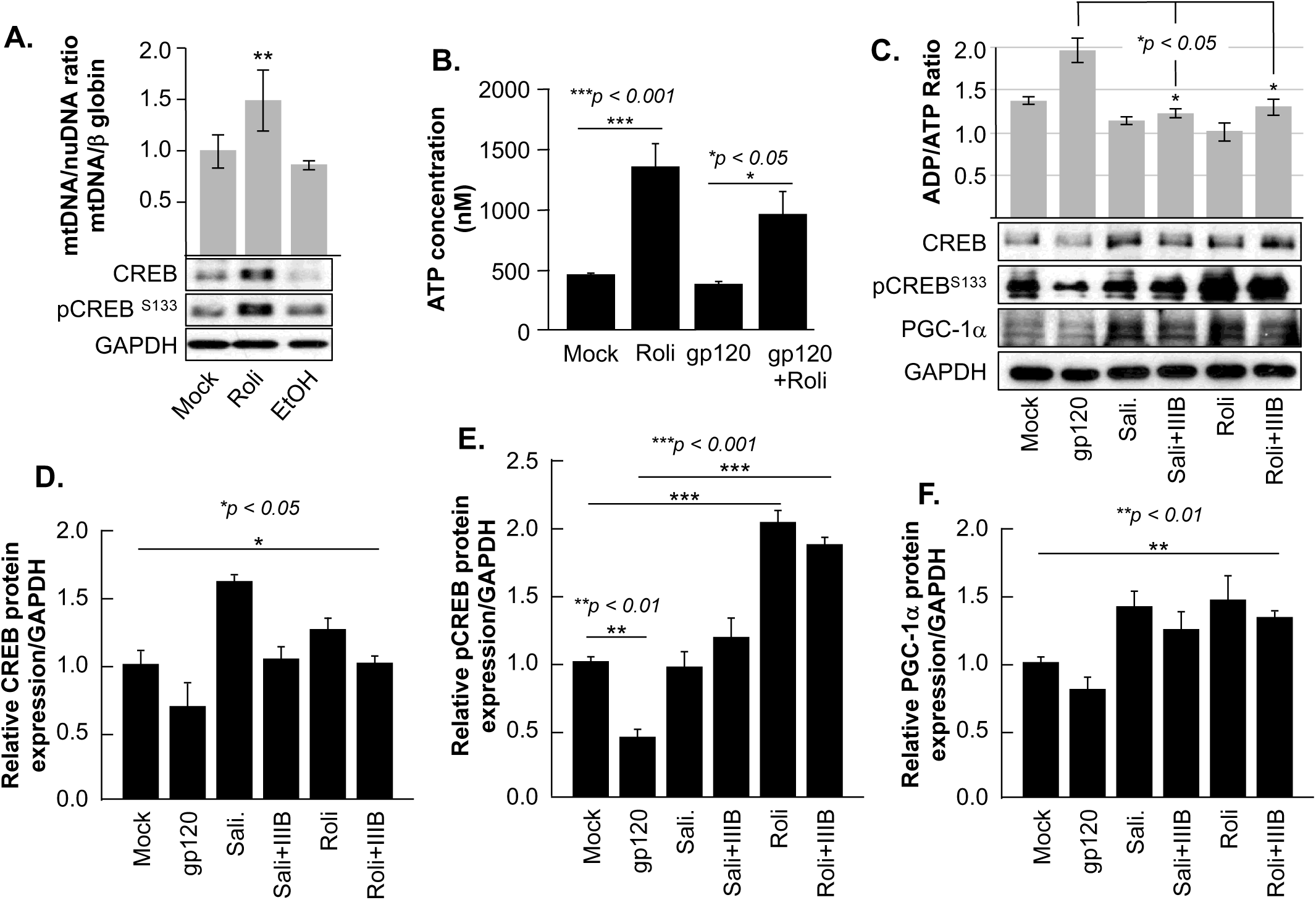
Rolipram restores CREB phosphorylation and targets. ***A***. HEK293 cells were treated with ethanol (200 mM for 24 hours) to repress and prevent pCREB^S133^ function or with rolipram (30μM/1hr) to induce pCREB^S133^ expression and function. mtDNA copy number represented by measuring the ratio of mtDNA/β-Globin. Expression levels of total CREB and pCREB^S133^ were also evaluated in HEK293 extracts using western blot analysis. Fold change in CREB expression relative to GAPDH was also measure and deemed significant (*p*<0.05). ***B***. Differentiated SH-SY5Y cells were treated with 30 μM of rolipram for 1 hour before adding 100 ng/ml of HIV-1 gp120 protein for 24 hours after which the reaction was stopped and cells collected. ATP levels in nM range were measured through a luciferase assay. ***C***. Measurement of ADP/ATP ratio in mock, HIV-1 gp120-treated, rolipram (30μM/1hr) ± HIV-1 gp120-treated, and salidroside (5μM/24hrs) ± HIV-1 gp120-treated SH-SY5Y cells as indicated. The ratio is presented as a bar graph and changes are considered significant (*p*<0.05) (n=3). *Lower panel* represents the protein expression of total CREB, pCREB^S133^, and PGC-1α in SH-SY5Y cells subjected to different treatment conditions as obtained by Western blot analysis. GAPDH is represented as the loading control. ***D, E, and F***. Quantitative analysis of the western blots presented along with the bands. The experiments were done in triplicate and the *p*-values were calculated using Student *t*-test (Mean ± S.D.). (\**p*<0.05; \*\**p*<0.01; \*\*\**p*<0.001).

As expected, EtOH treatment alone decreased mtDNA copy number (Fig 6A-histogram) and pCREB^S133^ expression level (Fig 6A, lower panel) while the addition of rolipram restored both (*the concentration and time of treatment of EtOH and Rolipram were chosen from publications using similar cells*).

Next, we measured the ATP production in the presence of rolipram. Differentiated SH-SY5Y cells were treated with rolipram (30 μM) for 1 hour and then treated with 100 ng/ml of recombinant HIV-1 gp120 for 24 hours before stopping the experiment. There was a decrease in the ATP level in the treated cells as compared to the untreated controls (Fig 6B). Further, while HIV-1 gp120 decreased ATP production, the addition of rolipram restored ATP levels.

Again, the ADP/ATP ratio was also measured in differentiated SH-SY5Y cells and treated with rolipram (30 μM) for 1 hour or with salidroside (5 μM) for 24 hours (another CREB activator) (46) before treating with HIV-1 gp120 for 24 hours (Fig 6C).

Here there is an increase in the ADP/ATP ratio in the presence of HIV-1 gp120 compared to the untreated control cells. The ratio decreased in cells treated with gp120 and rolipram and/or salidroside indicating more ATP production in the presence of these CREB activators (Fig 6C-Histogram).

Similarly, the addition of HIV-1 gp120 decreased the protein expression levels of total CREB, pCREB^S133^, and PGC-1α as shown by Western analysis (Fig 6C-lower panel). Interestingly, HIV-1 gp120 protein failed to lower expression levels of these proteins in the presence of rolipram or salidroside. Changes in CREB, pCREB^S133^, and PGC-1α protein expression levels were quantified using Image J and considered significant (Figs 6D, E, and F).

Our data presented in Figure 4D and E showed that the addition of HIV-1 gp120 protein affects mitochondrial movement and that could possibly be due to the loss of CREB’s ability to regulate PGC-1α. Since we have shown that rolipram can restore the expression level of PGC-1α (Figure 6C), we sought to determine whether the mitochondria movement can be restored as well. SH-SY5Y cells were transfected with 0.5 μg of Mito dsRed plasmid for 24 hours, differentiated, and then treated with rolipram (30 μM) for 1 hour before adding 100 ng/ml of HIV-1 gp120 for 24 hours. Moving mitochondria were observed using confocal microscopy and represented as a kymograph (Fig 7A). Treatment with HIV-1 gp120 protein reduced the ratio of mobile mitochondria from 38% to 18% (Fig 7B). There was a significant difference in the ratio of moving mitochondria in neurons treated with rolipram (59%) and/or rolipram + gp120 (50%) compared to the untreated or to HIV-1 gp120 treated cells excluding the possible false-positive effect brought by protein addition. The kymograph depicts the mitochondria movement in 5 minutes from left to right (anterograde transport). The average velocity of mitochondrial movement is presented in panel C.

**Figure 7.**
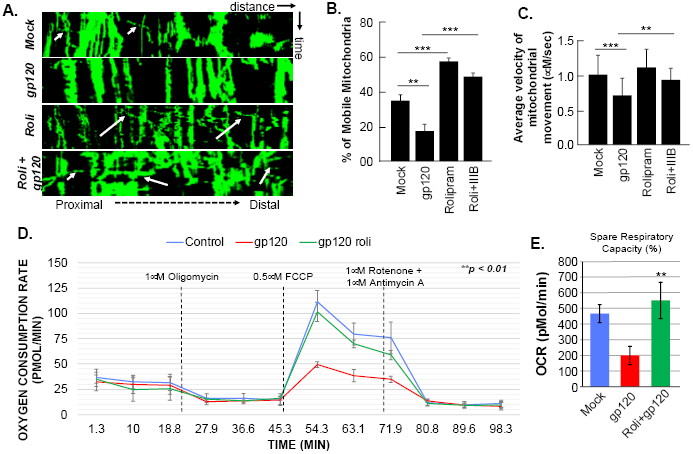
Rolipram treatment improves mitochondrial movement and oxygen consumption rate. ***A***. Kymographs representing the mitochondrial movement in mock, HIV-1 gp120-treated (100ng/ml), rolipram-treated, and rolipram + HIV-1 gp120-treated SH-SY5Y cells. Horizontal axis represents the distance and vertical axis represents the time. Mobile mitochondria are distinguished by the diagonal dotted lines moving along the horizontal axis (arrows) and stationary mitochondria are represented by vertical lines. The mitochondria were tagged using Mito ds-RED transfection and the movement was visualized using Leica EL600 DMI3000 confocal microscopy. ImageJ was used to create a kymograph. Data is the representation of at least 15 - 20 neurite per treatment condition each repeated 3 times (n= 60 - 65 total neurites per treatment conditions). ***B***. Quantification of the percentage of mobile mitochondria. ***C***. Average velocity of the mobile mitochondria (n = 60 - 65 per treatment condition). Data represents the mean ± S.D. Results were judged statistically significant if *p*<0.05 by analysis of variance. (\**p*<0.05; \*\**p*<0.01; \*\*\**p*<0.001). ***D***. Differentiated LUHMES cells (500 cells/μl) were treated with 100 ng/ml of HIV-1 gp120 protein for 24 hours after which oxygen consumption rate (OCR) and extracellular acidification rate (ECAR) were measured using the Agilent Seahorse Mitochondrial stress test as displayed. Measurements were made at basal conditions, and in response to 1 μM Oligomycin, 1.5 μM FCCP, and 1 μM rotenone + 1 μM antimycin A. Experiments were performed in triplicate for each experimental group and considered significant. ***E***. OCR of cells represented as percent spare respiratory capacity (difference between OCR immediately before addition rotenone and antimycin A and OCR after). Control/mock represented in blue, HIV-1 gp120-treated cells in red, and rolipram + HIV-1 gp120-treated cells in green. Data represents the mean ± S.D. Results were judged statistically significant (\*\**p*<0.01).

To further validate our hypothesis and to really look at mitochondrial function, we measured the mitochondrial oxygen consumption rate via Mito Stress Seahorse Assay (Fig 7D). Luhmes cells were differentiated and grown on Agilent Seahorse XFe24 well plates and treated with 100 ng/ml HIV-1 gp120 for 24 hours. The oxygen consumption rate (OCR) of the cells was then measured in response to various mitochondrial-stressing agents. The addition of HIV-1 gp120 decreased OCR in response to FCCP (0.5 μM), an uncoupler that triggers the respiratory chain and simulates maximum capacity, suggesting that these cells do not have the ability to perform in an energy-demanding state. In the presence of rolipram, however, the OCR is restored. Similarly, the percentage of spare respiration capacity, another indicator of cell fitness, decreased in the presence of HIV-1 gp120 but was restored when rolipram was added (Fig 7E).

Interestingly, all measurements of mitochondrial dysfunction caused by the addition of HIV-1 gp120 protein were reversed in the presence of rolipram or salidroside. These results indicate a rescue effect of treatment with CREB activators, thus indicating the role of CREB in HIV-1 gp120-induced mitochondrial dysfunction.

### Evidence that Rolipram restored memory in mice

Our data show rolipram can restore CREB functions *in vitro* so here we sought to determine its effect *in vivo* using a mouse model. C57Bl/6J mice received stereotaxic injections of gp120 (125ng/μl) or saline bilaterally into the hippocampi, followed by IP injection of rolipram (1 mg/kg) or saline immediately after surgery and 36 hours after surgery. 72 hours post stereotaxic injection, mice underwent object location memory training. After a 24-hour delay, spatial memory was tested (Fig 8A).

**Fig 8.**
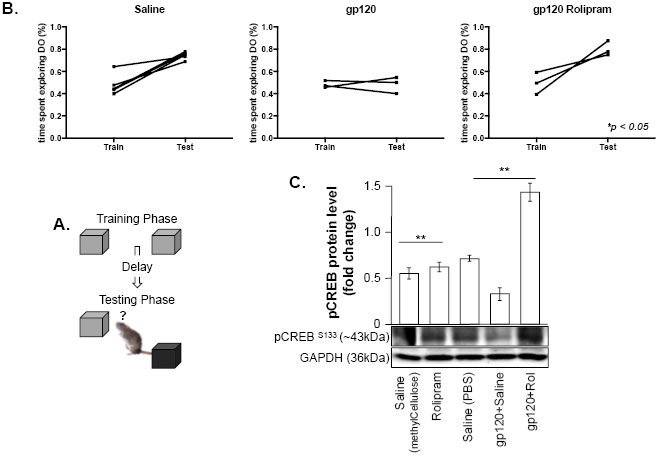
Restoring learning and short memory with rolipram. ***A***. Schematic representation of the object location spatial memory test. ***B***. Mice were injected with either saline (left), gp120 protein followed by saline (middle), or gp120 followed by rolipram (1mg/kg, right). 72 hours later, mice were trained and tested in the object location spatial memory task. The amount of time spent exploring the object to be displaced was recorded during the training phase, and the amount of time spent exploring the displaced object during the test phase was recorded. Time spent with the displaced objects during the test phase compared to the training phase was assessed using two tail *t*-test (*p*<0.05) as indicated. ***C***. Proteins were extracted from the brain tissues (hippocampal area) of saline, gp120, and/or gp120/rolipram-injected mice and subjected to Western blot as indicated. Data represents the mean ± S.D. Results were judged statistically significant (\*\**p*<0.01).

Mice that received stereotaxic injections of saline control exhibited normal long-term spatial memory (measured as time spent exploring), whereas mice that received HIV-1 gp120 injections had impaired memory as expected (Fig 8B). However, we found that the injection of rolipram in the presence of HIV-1 gp120 restored memory. Donepezil, a CREB inducer, also restored memory (data not shown) (47). Brains were collected from these mice and protein extracts were isolated from the hippocampal area and then subjected to western blot analysis. As shown, the expression of phosphorylated CREB was decreased in HIV-1 gp120-injected mice but maintained following rolipram injection compared to saline-injected mice (Fig 8C). These data suggest HIV-1 gp120-induced CREB deregulation and subsequent mitochondrial dysfunction plays a role in memory impairment *in vivo*.

## DISCUSSION

HIV-1 enters the brain early in the course of infection and begins to evolve separately from the blood-based viral population. Available evidence suggests that HIV-associated neurocognitive disorder (HAND) is an indirect consequence of HIV-1 infection where viral proteins (e.g. gp120) and inflammatory mediators released by infected cells damage the neurons. Typical features of HAND include spatial memory loss, inability to manipulate acquired knowledge, and generalized slowing of thought processes that are similar to symptoms observed in older yet overall healthy people. To examine these features, we showed the impact of HIV-1 gp120 protein on spatial memory impairment *in vitro* and *in vivo*. Looking for the mechanisms involved, we demonstrated the ability of HIV-1 gp120 to decrease CREB protein expression (total and phosphorylated) and function.

The role of CREB in spatial memory formation and synaptic plasticity has been amply studied. However, these features have never been explored in the context of HAND. Nevertheless, CREB’s functional relation with HIV-1 proteins has been moderately examined. CREB was shown to bind and to regulate the HIV-1 promoter activity in HeLa and Jurkat cells (48). It was also demonstrated to functionally interact with HIV-Tat protein. HIV-1 Tat has been shown to utilize CREB to promote IL-10 production, though the significance of this functional interaction regarding HIV pathogenesis remains unclear (49). Further, several studies showed a decreased expression of CREB in neurons treated with HIV-Tat and/or gp120 proteins, however, CREB has never been the subject of these studies, and the mechanisms involved were not examined (50, 51). Hence, this current study presents a novel hypothesis and it clarifies the relationship between gp120 and CREB.

In this study, we have shown that HIV-1 gp120 protein causes the loss of CREB phosphorylation and function promoting mitochondrial perturbation such as inhibition of PGC-1α, the disappearance of mitochondrial cristae, and decreased ATP production (Fig 4). While a functional interaction between CREB and PGC-1α has been described, the relationship between HIV-1 gp120 and PGC-1α has never been examined and this is the first study to demonstrate such a relation. CREB has been shown to regulate the function of PGC-1α directly by binding to its promoter or indirectly by lifting its methylation using CREB’s downstream target, miR-132 (52). Being a downstream target of CREB, using SH-SY5Y cells, we demonstrated that by lowering the expression level of CREB protein, gp120 also decreased miR-132 expression and function (data not shown). Loss of miR-132 activates DNMT3b which in turn alters the PGC-1α promoter (52). Interestingly, in astrocytes, HIV-1 Tat was shown to activates miR-132 expression which diminishes the MeCP2 level and negatively affects neurite length (53). Based on these results, we concluded that a functional feedback loop exists between HIV-1 gp120, PGC-1α, and CREB (Fig 3B).

In addition to PGC-1α, here we demonstrated that HIV-1 gp120 also caused a decrease in BDNF protein expression and shortening of the neurites, both involved in synaptic plasticity and memory function (Fig 5). Several groups have already shown the negative impact of HIV-1 gp120 on BDNF protein expression and function (54). Most of these studies focused on the role of the p75 neurotrophin receptor and linked the effect of HIV-1 gp120 on BDNF to this receptor (55). We believe that these results are lacking the fact that HIV-1 gp120 alters BDNF function by preventing CREB from binding to the CRE site with the BDNF promoter IV (data not shown).

Our data suggest that HIV-1 gp120 protein impairs spatial memory through two major pathways. The first one (Tricarboxylic acid cycle – TCA, aka Krebs cycle), described in this study where HIV-1 gp120 can lower CREB function leading to mitochondria and synaptic plasticity deregulation through PGC-1α and BDNF (low ATP, disrupted NAD/NADH, loss of mitochondria movement, and neurite retraction). While the second pathway (glycolysis) involved different players that could affect CREB activity indirectly.

A decrease in ATP and NADH levels could also mean that gp120 is favoring the glycolysis pathway as opposed to the regular TCA pathway. In this regard, one may ask, if the NAD/NADH ratio is disrupted, how the cell can survive and remain functional. It has been shown that when/if the glycolysis pathway is activated, the cell will use its tryptophan which is a part of the kynurenine (KYN) pathway, this phenomenon is well described in cancer (56). KYN pathway has also been shown to be linked to depression and cognitive impairments in HIV-1 patients (57). Interestingly, the conversion of KYN to kynurenine acid (KYNA) is somehow under the control of PGC-1α, and the loss of PGC-1α promotes the KYN toxicity (58). This provides additional evidence that gp120 is causing memory impairment through promoting the loss of CREB function that could affect its downstream targets such as PGC-1α which could increase inflammation and depression. Note that the *pgc-1*α gene can produce several isoforms which can play different roles and respond to viral proteins differently (59). This might explain the different behavior of HIV-1 clade B and C. Further, the glycolysis pathway leads to the activation of the inducible cAMP early repressor (ICER) that prevents CREB protein from interacting with its specific DNA binding site (60), which in our case could be the CRE sites located within the CREB, PGC-1α, and/or BDNF promoters, hence leading to spatial memory impairment.

The use of rolipram, salidroside, or donepezil shed the light on an additional understudied pathway, the one that involves phosphodiesterase protein (PDE4). CREB loss of phosphorylation can be the result of cAMP degradation by PDE4, which prevents CREB phosphorylation and binding to its cognate site. The addition of rolipram which is a phosphodiesterase antagonist permits the phosphorylation of CREB, neutralizes the gp120 effect, and restores memory function as observed in Figures 6 and 7. This observation raises the possibility that gp120 could cause CREB loss of function and impaired spatial memory through activation of PDE4, however, this has yet to be explored. The relation between PDE4 and HIV-1 was described in one study where the authors showed that selective blocking of PDE4 activity inhibits IL-2R expression and thereby leads to abolishing HIV-1 DNA nuclear import in memory T cells (61). Therefore, to design a therapy against spatial memory impairment, our future studies should focus on exploring the additional two pathways (glycolysis and PDE4).

To further validate the impact of gp120 on memory impairment, we performed proteomics studies where we showed that the addition of gp120 protein, clade B but not clade C, negatively regulates the CREB and Synaptogenesis pathways in SH-SY5Y cells (data not shown), which confirm that gp120B negative effect on memory is specific and depends on CREB pathway.

In summary, we presented evidence that circulating gp120 protein, which could be secreted by defective proviruses, has the capability of altering the CREB pathway and CREB downstream targets such as PGC-1α and BDNF. All of which can lead to the loss of spatial memory. Understanding these mechanisms can help to develop better therapy to prevent spatial memory loss observed in a significant number of HIV-infected patients. Further, deciphering these mechanisms could also help in understanding the development of other neurodegenerative diseases.

## EXPERIMENTAL PROCEDURES

### Cell culture and Treatments

The human neuroblastoma cell line, SH-SY5Y, was purchased from ATCC (CRL-2266) and grown in Dulbecco modified eagle medium: nutrient type - 12 (DMEM F12) supplemented with 10% fetal bovine serum (FBS), 1% non - essential amino acid and 1% sodium pyruvate. All cells were incubated at 37°C supplemented with 5% CO_2_. The cells were passed each time at 85 - 90% confluency. The cells only within 10 passages from the time of purchase were used. The cells were seeded at the density of 5 × 10^5^ cells/per well in 6 well plates. They were differentiated into neurons with 10 μM retinoic acid (RA) treatment for at least 3-4 days.

HEK293T cells were purchased from ATCC and grown in DMEM supplemented with 10% FBS. All cells were incubated at 37°C supplemented with 5% CO_2_. The cells were passed each time at 85 - 90% confluency. The cells were seeded at the density of 5 × 10^5^ cells/per well in 6 well plates..

LUHMES cells were purchased from ATCC, all cell culture dishes are pre-coated with 50 μg/ml PLO and 1μg/ml fibronectin and incubated at 37°C for 3 hours. All cells were grown in DMEM/F12 with 1× N2 supplement, 2 mM L-glutamine, and 40 ng/ml recombinant bFGF. LUHMES are then differentiated for 6 days in complete differentiation media (DMEM/F12, 1× N2 supplement, 2mM L-glutamine, 1μg/ml doxycycline, and 2ng/ml recombinant human GDNF).

Human primary neuronal cells (26 weeks old fetal brain) were generously provided by Dr. Elyseo Eugenin from Rutger’s University (*Current address: UTMB*). The human cortical fetal tissue was used to isolate the mix culture of neurons and astrocytes as described (62). The culture obtained contained 30-40% neurons, 60-70% astrocytes and 2-5% microglia. Neuronal enriched cultures were obtained after 7-10 days of culture in Neurobasal medium supplemented with N2 neuro-survival factor and 5% FBS. The cells were cultured for about 10 days to obtain 70-80% of the neuronal culture before the treatments. (*note that human primary cultures experiments were performed prior to 2016*).

Primary C57B1/6J mouse neurons were obtained from E18 (embryonic day 18). The cortex and hippocampus were separated and rinsed in HBSS before digesting in 0.125% trypsin. The cortex and hippocampus were digested for 15 and 20 minutes, respectively (DNase 0.4%). The digested tissue was triturated in DMEM containing 10% FBS followed by centrifugation at 1000 rpm for 10 minutes. The tissue was then washed twice, re-suspended in DMEM F12 and passed through a strainer to remove the non-dispersed tissue. The cells were seeded at 7 × 10^5^ cells/well in 6-well plates and 1.2 × 10^5^ cells/chamber in 4-chamber slides and incubated at 37°C supplemented with 5% CO_2_. DMEM medium was replaced with Neurobasal supplemented with B27, NEAA, GluMax and pen-strep after 2-3 hours of seeding. Half a medium was changed every 3 days.

### HIV-1 gp120 Treatments

Recombinant HIV-1_IIIB_ gp120 (clade B) and HIV-1_CSF_ gp120 (clade C) proteins were kindly received from NIH AIDS Reagent Program (63). Samples were treated for 24 or 48 hours in concentration indicated. HIV-1_IIIB_ gp120 is T-tropic and works via CXCR4 receptor.

### Chemical Reagents

AMD3100 (gp120-CXCR4 inhibitor) was kindly received from NIH AIDS Research Program and used at a concentration of 1μM. Cells were treated with rolipram (30 μM, purchased from Sigma-Aldrich) or with salidroside (5 μM, purchased from Cayman Chemicals) for 1 and 24 hours, respectively prior to the addition of gp120 protein. Both chemicals are described as activators of cAMP. Donepezil was used at a concentration 1mg/kg.

### Cell viability assay (MTT assay)

The cell viability of control and gp120-treated and differentiated SH-SY5Y cells was measured using a standard MTT assay. Briefly, 5 × 10^5^ viable cells were seeded in triplicate into clear 96-well flat-bottom plates in media supplemented with 10% fetal bovine serum and incubated with different concentrations of the gp120 protein for 24 and 48 hours. Then, 10 μl/well of MTT (5 mg/ml) was added and the cells were incubated for 4 h. Following incubation, 100 μl of 10% sodium dodecyl sulfate (SDS) solution in deionized water was added to each well and left overnight. The absorbance was measured at 595 nm in a benchtop multimode reader (Molecular Device).

### Western Blot assay

Proteins were extracted using radioimmunoprecipitation assay (RIPA) lysis buffer (25 mM Tris–HCl pH 7.6, 150 mM NaCl, 1 % Triton and 0.1 % SDS) + protease inhibitor cocktail. Protein concentrations were estimated using Bradford Assay (Bio-Rad). Western blot was performed using 25μg of extracts per well and anti-CREB, - p^S133^CREB, -PGC-1α, -Miro, -BDNF (N20), -PSD-95, (Santa Cruz), -TRAK1, -KIF1β, -Dynein, (Abcam), and -GAPDH antibodies. Densitometry ratio of the bands were determined using an ImageJ that was normalized to the GAPDH. GAPDH was used for equal proteins loading.

### RNA extraction

Total RNA was extracted from the sample using SurePrep TrueTotal RNA purification kit from Fisher Bioreagents. Nanodrop was used to determine the purity and concentration of the RNA extracted.

### qPCR assay

cDNA was synthesized using SuperScript VILO cDNA synthesis kit (Invitrogen). The following primers were purchased from IDT: BDNF: (F)-5’-gagcagctgccttgatggttactt-3’; (R)-5’-a agccaccttgtcct cggatgttt-3’. CREB: (F)- 5’-ggcagacagttcaagtccatg-3’; (R)- 5’-cgctttgggaatcagttacac-3’. PGC-1α: (F)- 5’-ccaaaccaacaactttatctc-3’; (R)- 5’-cacttaaggtgcgttcaa tag-3’. GAPDH: (F)- 5’ gccttccgtgttcctacc-3’; (R)-5’-cctcagtgtagcccaagatg-3’. Results are expressed in relative gene expression level as compared to the untreated control. GAPDH was used as an internal control.

### Immunohistochemistry

#### a- Human brain

Frontal lobe brain tissues from HIV-positive patients with varying degrees of dementia, along with non-demented and HIV-negative controls were obtained from the national neuroAIDS tissue consortium (NNTC). The formalin-fixed and paraffin-embedded tissues were sectioned at 5μm thickness and placed on electromagnetically charged glass slides. Sections were deparaffinized in xylene and re-hydrated through descending grades of alcohol up to water. Non-enzymatic antigen retrieval was performed in citrate buffer for 30 minutes at 95°C in a vacuum oven. Sections were then rinsed with PBS and permeabilized in 0.2% Triton in PBS for 45 minutes at room temperature. Sections were rinsed again with PBS, and a blocking step was performed with normal BSA serum at room temperature in a humidified chamber for 2 hours Primary antibodies were incubated overnight at 4°C and later for 1 hour at room temperature with fluorescently labeled secondary antibodies. The tissues were subsequently washed in PBS until finally mounted with DAPI containing medium (Vectashield). Leica EL600 DMI3000 confocal microscope was used for imaging.

#### b- gp120 transgenic mice

Imaging of gp120-tg mouse brains was done in Dr Kaul’s lab (Sanford Burnham Presbys Medical Discovery Institute). WT and gp120-tg mice (express gp120 under GFAP promoter) were kindly provided by Dr Lennart Mucke (Gladstone Institute of Neurological Disease, University of California) (30) and maintained at Sanford Burnham Presbys Medical Discovery Institute. The mice (WT and gp120-tg), 9 months of age, were anesthetized with Isoflurane and transcardially perfused with 0.9% saline. The brains were quickly removed and fixed with 4% paraformaldehyde for 48 hours at 4°C. The brain sections of 30 μm thickness were obtained for the histological studies. The slides were permeabilized with 1% Triton X-100 for 30 mins followed by blocking with 10% heat/inactivated goat serum in PBS containing 0.5% Tween 20 for 1.5 hours. The sections were then stained with CREB (Cell Signaling) and MAP-2 (Sigma) overnight followed by Alexa Flour 488-labled goat anti-rabbit (Molecular Probes). Nuclear DNA was labeled with H33342. Per animal at least three sagittal sections were analyzed and each sections five fields were recorded using Zeiss inverted Axiovert 100M fluorescence microscope (29). All experiments involving gp120tg mice were performed in accordance with NIH guidelines and approved by the IACUC of Sanford Burnham Presbys Medical Discovery Institute.

### mtDNA copy number

Total genomic DNA was extracted from differentiated and gp120 treated SH-SY5Y cells with different treatment conditions. To measure mtDNA quantitative real time PCR was performed with SYBR green (ROCHE). The primer sequences used for mtDNA were: mt (F)- 5’-cgaaaggacaagagaaataa gg-3’ and mt (R)- 5’-ctgtaaagttttaagttttatgcg-3’. β-Globin (F)- 5’-caacttcatccacgttcacc-3’ and (R)- 5’-gaagagccaaggacaggtac-3’. The primers were purchased from Integrated DNA Technologies (IDT-Illinois-USA). Note that mtDNA was measured using qPCR while the copy number was measured as a ratio of mt gene from the D loop over a nuclear gene (β globin).

### Mitochondrial Mobility (live cell images)

SH-SY5Y cells were seeded on the glass bottom plates and transfected with 0.5 μg of Mito dsRED plasmid (Clontech) using lipofectamine 2000 (Invitrogen). The transfection media (Opti-MEM) was replaced with the differentiation media after 4-6 hours and the cells were left to differentiate for 96 hours and then treated with 100 ng/ml of gp120 protein for 24 hours and/or 30. μM of rolipram for 1 hour. The cells were visualized using Leica confocal microscope Leica (DMI4000) for live cell imaging. A heated 37°C temperature-controlling chamber filled with 5% CO_2_ surrounding the microscope stage was used to keep the cells alive. Images were captured every 5 seconds for a total of 5 minutes. The area of the cell visualized was focused on the processes and not the cell body. Mitochondrial movement along the processes were represented in the form of kymograph by using Image J. Mitochondria moving at a velocity higher than 0.1 μm/s were considered mobile. At least thirty axons were recorded. The vertical straight line represents the immobile mitochondria whereas, the dotted line along the horizontal axis represents the mobile mitochondria.

### ATP assay

ATP was measured with a luciferin – luciferase bioluminescence assay using ATP determination kit from Molecular Probes as per manufacturer’s protocol. Briefly, the standard curve was measured using different concentration of ATP solution (1nM - 1μM) with the standard solution. The cells were subjected to freeze and thaw cycle (3×) to release the ATP. The cells were centrifuged at 12000 rpm for 10 minutes. 10μl of supernatant was used for every 100μl of the standard solution.

### ADP/ATP ratio

Changes in the ADP/ATP ratio were measured using the ADP/ATP ratio assay kit purchased from Sigma-Aldrich which is based on a luciferin - luciferase assay. The assay involved two steps: (i) ATP reacts with the substrate D luciferin and produce light in the presence of luciferase - measuring the intracellular ATP concentration; (ii) ADP is converted to ATP through an enzymatic reaction, followed by the reaction of ATP with D-luciferin. The second light intensity measurement represents the total ADP and ATP concentration in the sample. The ATP reagents were prepared where 90μl/well were added to the samples and incubated for 1 minute at room temperature followed by luminescence reading. This operation was repeated with 10 minutes incubation. Immediately after the second reading 5 μl of ADP reagent was added to each well, incubated for 1 minute at RT, and the luminescence was read. The ADP/ATP ratio was then calculated.

### NAD^+^/NADH Ratio

Differentiated and gp120-treated SH-SY5Y cells were washed with cold PBS, and centrifuged at 2,000 rpm for 5 minutes. The cells were homogenized with 400 μl of NADH/NAD buffer, vortexed for 10 seconds and then centrifuged at 13,000*g* for 10 minutes to remove any insoluble materials. The extracted NAD/NADH supernatants were then transferred into a 96-well plate with a final volume of 50 μl to measure NAD^+^. The second set was prepared by aliquoting 200 μl of the samples into microcentrifuge tubes and heating them to 60°C for 30 minutes. The samples were then cooled on ice, spun and transferred into the plate. This is used to measure NADH. 100μl of the Master Reaction and 10 μl of the NADH Developer were added to each sample. The absorbance was measured at 450nm at different time points (30, 60 and 120 minutes) using a Modulus microplate reader. The ratio of NAD^+^/NADH was calculated.

### ROS assay

The trafficking of 2,3,4,5,6-pentafluorodihydro-tetramethylfosamine (redox sensor red CC-1; Invitrogen) was used to detect reactive oxygen intermediates. Redox Sensor Red CC-1 is oxidized in the cytosol and accumulates in either the mitochondria or the lysosomes depending on the oxidation of the cell. Thus, Redox Sensor Red is a measurement of redox potential. Differentiated SH-SY5Y cells under different treatment conditions were incubated for 5 minutes with 1μm of Redox Sensor Red CC-1 and a mitochondria-specific dye, MitoTracker Green FM (25 nm; Molecular Probes). 100 nM H_2_O_2_ was used as a positive control. Culture slides were washed with 1X PBS and visualized using an EVOS cell imaging system. If Redox Sensor Red is co-localized with MitoTracker Green in the mitochondria then the oxidation level of the cell is high. Note that all ROS measurement experiments were performed at least three times.

### Mitochondria stress test

100 μl of LUHMES cells on day 5 of differentiation were plated onto 50 μg/ml PLO and 1μg/ml fibronectin coated Agilent Seahorse XFe24 well plates at a density of 500 cells/μl. After 24 hours the cells were treated with 100ng of HIV-1 gp120 protein and allowed to incubate for another 24 hours. After which oxygen consumption rate (OCR) and extracellular acidification rate (ECAR) were measured using the Agilent Seahorse Mitochondrial stress test. The cell media was changed to the XF media (DMEM,10 mM glucose, 4 mM L-Glutamine, and 2 mM sodium pyruvate). Using the XF24 Seahorse analyzer, ECAR and OCR measurements were made at basal conditions, and in response to 1 μM Oligomycin, 1.5 μM FCCP, and 1 μM rotenone + 1μM antimycin A. Experiments were performed in triplicate for each experimental group.

### Synaptophysin vesicles number and Neurite retraction and distribution assays

CellLight Synaptophysin RFP BacMac 2.0 from Life Technology (catalog# C10610) was used to transduce differentiated primary mice neurons or SH-SY5Y cells at MOI of 5 for 24 hours following the manufacturer’s protocol. Images were taken using the EVOS microscope and Image J was used to count the number of synaptophysin vesicles or to measure neurite retraction.

### Transmission Electron microscopy (TEM)

Differentiated SH-SY5Y cells were grown on 100 × 20 mm tissue culture treated dishes (Celltreat^®^ Scientific Products, MA) for 3 days in F-12/DMEM (50/50) supplemented with sodium pyruvate, nonessential amino acids, and 10% FBS. Cells were then treated with 100 ng/mL gp120 recombinant Protein or PBS for 24 hours. The cells were then collected, centrifuged and fixed. To fix the cell, 500μl of formaldehyde glutaraldehyde, 2.5% in 0.1 M Sod. Cac. buffer, pH 7.4 (Electron Microscopy Sciences, Hatfield, PA) was added slowly to the top of the pellet. The samples were then put on ice and transferred to the EM facility. After fixation cells were centrifuged at 1000g, washed 3 times with 0.1 M Cacodylate buffer. The cell pellet (~ 50 μl) was mixed with 100 μl of 5% agarose and centrifuged at 1000*g* for 10 minutes. Tubes with cell pellets were transferred at 4°C/ice for 1 hour to solidify agarose. The cell pellets in agarose were post fixed with 1% OsO_4_ in solution for 1 hour at room temperature and washed 3 times with DD H_2_O. Samples were further processed for dehydration with serial changes of Ethanol, were embedded into Epon 812 resin (Electron Microscope Sciences, USA) and polymerized at 65°C for 72 hours. Embedded samples were cut thin (80-100 nm) using Leica UC6 microtome, collected on 200 mesh copper grids (Electron Microscopy Sciences, USA) and stained with 2% Uranyl Acetate for 12 min at RT followed by staining with Reynolds lead citrate for 6 minutes at RT. Samples were visualized with FEI Tecnai T12 transmission electron microscope, at 100kV, equipped with 2K x 2K Megaplus camera Model ES 4.0 (Roper Scientific MASD, San Diego CA). Mitochondria are highlighted in blue and numbered before using image J to measure area and perimeter. Any structure that cannot be clearly identified as a mitochondrion (i.e. no evidence of cristae or double membrane) or that has been cut off when acquiring the image was not included. Standard error was used (p=0.378 [*t*-test]).

### Stereotaxic surgery and spatial memory testing

#### Stereotaxic surgery

Mice were singly housed with *ad-libitum* access to food and water prior to and after surgery. 8-10 weeks old C57B1/6J male mice were anesthetized with isoflurane and received bilateral stereotaxic injections (1 μL volume per injection site) of either saline or gp120 (125 μg/mL) into the hippocampi at rostral (−1.7 mm A/P, 1.2 mm M/L, 2 mm D/V from bregma) and caudal (−2.7 mm A/P, 2 mm M/L, 2.1 mm D/V from bregma) coordinates for a total of 4 intrahippocampal injections per mouse. Mice then received intraperitoneal injections of saline or rolipram (1mg/kg) immediately after surgery and 36 hours after surgery. 72 hours post stereotaxic injection, spatial memory was tested using the object location memory test. All procedures were approved by the Baylor College of Medicine Institutional Animal Care and Use Committee.

#### Object location memory testing

The object location memory test requires that mice learn and remember the positions of two objects in an arena. Extra-arena spatial cues exist to orient the mice during training and testing phases. For training, two identical flasks were placed at adjacent far corners of the arena, and animals were allowed to explore both flasks in 3 trials of 3 min each with 3 min intervals. The amount of time mice spent exploring each flask was recorded by the experimenter. After a delay of 24 hours, mice were returned to the arena for the test phase. In this phase, one flask was displaced to the adjacent empty corner. Mice were given 3 minutes to explore both flasks, and the amount of time spent exploring each flask was recorded. An increase in time spent with the displaced object during the test phase relative to the training phase is indicative of intact spatial memory.

### Statistical Analysis

All the experiments were repeated at least in triplicate. Statistical analysis was performed using one-way analysis of variance with a *post hoc* Student’s *t*-test. Data are expressed as the mean of ± S.D. Results were judged statistically significant if *p*<0.05 by analysis of variance. (Marked in the figure as \**p*<0.05; \*\**p*<0.01; \*\*\**p*<0.001 where needed). Data were plotted either using GraphPad Prism version 5.0 or 7.0.

## DATA AVAILABILITY

All processed data are included in this manuscript. Raw data, further information, or reagents contained within the manuscript are available upon request from the corresponding author, Bassel E Sawaya, Lewis Katz School of Medicine - Temple University Philadelphia, PA.

## ACKNOWLEDGMENTS

This work is supported by an NIH-NIA grant AG054411 awarded to BES, NIH-NINDS grants NS085171 and NS086965 awarded to JC, and NIH grants MH087332, MH104131, MH105330 and P50 DA026306 (P5) to MK.

The following reagents were obtained through the NIH AIDS Reagent Program, Division of AIDS, NIAID, NIH: HIV-1 IIIB gp120 Recombinant Protein from ImmunoDX, LLC; HIV-1 JR-CSF Fc-gp120 Recombinant Protein (Cat#11556) from Aymeric de Parseval and Dr. John H. Elder.

## AUTHOR CONTRIBUTIONS

JS, MS,CAN, SPA, AB and MK conceived the study. RM shared reagents. JP performed animal studies. JS, JC and BES analyzed experiments. JS, SPA and BES prepared figures and wrote manuscript. BES planned and directed the work. The manuscript was edited by all authors.

## ABBREVIATIONS

HIV: Human immunodeficiency virus
HAND: HIV-associated neurocognitive disorders
CSF: cerebrospinal fluid
cART: combinatory antiretroviral therapy
CREB: cyclic AMP response element-binding protein
PGC-1α: peroxisome proliferator-activated receptor gamma coactivator
BDNF: brain-derived neurotrophic factor
OS: oxidative stress
AD: Alzheimer’s disease
PD: Parkinson’s disease
ALS: Amyotrophic lateral sclerosis
HD: Huntington’s disease
CRE: cAMP response elements
IL-10: interleukin 10
PKA: protein kinase A
tg: transgenic
GFAP: glial fibrillary acidic protein
IF: immunofluorescence
ROS: reactive oxygen species
post synaptic density 95: PSD-95
mtDNA: mitochondrial DNA
OCR: oxygen consumption rate
ECAR: extracellular acidification rate
TCA: Tricarboxylic acid cycle

